# Functional, transcriptional, and microbial shifts associated with healthy pulmonary aging: insights from rhesus macaques

**DOI:** 10.1101/2022.02.08.479578

**Authors:** Nicholas S. Rhoades, Michael Davies, Sloan A. Lewis, Isaac R. Cinco, Steven G. Kohama, Luiz E. Bermudez, Kevin L. Winthrop, Cristina Fuss, Julie A. Mattison, Eliot R. Spindel, Ilhem Messaoudi

## Abstract

Older individuals are at increased risk of developing severe respiratory infections due to age-related changes in the immunological, microbial, and functional landscape of the lung. However, our understanding of the impact of age on the respiratory tract remains limited as samples from healthy humans are challenging to obtain and confounding variables such as smoking and environmental pollutant exposure make it difficult to assess the true impact of aging. On the other hand, studies in rodent models are biased by their specific pathogen free status. In this study, we utilize a rhesus macaque model of healthy aging to examine the functional, immunological, and microbial consequences of aging in the lung. Pulmonary function testing in this large (n=34 adult, n=49 aged) cross-sectional study established age and sex differences similar to humans supporting the translational accuracy of this model. Additionally, an increased abundance of myeloid cells (alveolar and infiltrating macrophages) and a concomitant decrease in T-cells were also observed in aged animals. Single cell RNA sequencing indicated a transcriptional shift in the pulmonary CD8+ T-cell population from *GRZMB* expressing cells to *IFN* expressing cells, while frequency of *IL-1B* expressing alveolar macrophages was significantly reduced. Interestingly, the lung microbiome of many animals was dominated by a single microbe, *Tropheryma* spp., the prevalence of which decreased with age. These data provide a comprehensive picture of the functional, microbial and immunological changes of the lung in healthy macaque aging and provide insight into the increased prevalence and severity of respiratory disease in the elderly.

## INTRODUCTION

The number of older individuals (>65 years old) is rapidly growing worldwide (1). Indeed, it is estimated that 16% of the world population will be older than 65 years of age (up from 9% in 2019) and that the proportion of the oldest old (>85 years of age) is expected to triple by 2050 (1). This shift in demographics presents significant challenges to the healthcare system as older individuals are more susceptible to microbial infections and more likely to develop severe disease necessitating hospitalizations and admission to the intensive care unit (2, 3). Aged persons are exponentially more vulnerable to respiratory diseases as highlighted by the fact that chronic lower respiratory disease are the third leading cause of death for aged individuals (4). Moreover, advanced age is associated with severe respiratory bacterial, fungal, and viral infections (5–7). For instance, death from influenza/pneumonia remains one of the leading causes of mortality in the elderly (2, 8). This pattern can be at least partially attributed to physiological and structural changes in the lungs that have implications for host health. For example, aging leads to impaired mucociliary clearance and cough strength (9–11) resulting in decreased ability to clear particles from the lung (11). Aging also leads to decreased expiratory flows, volumes and overall gas exchange capability (12–17). However, the relationship between these physiological changes and age-associated shifts in lung immunity and microbiome are poorly understood.

It is well documented that aging leads to drastic shifts in systemic immunity, notably a loss of naive T-cells that is accompanied by accumulation of terminally differentiated T-cells, and development of low-grade chronic inflammation referred to as inflammaging (18–20). These changes lead to reduced antimicrobial defenses and responses to vaccination (21–23). In contrast to age-related changes in peripheral immunity, changes at mucosal sites such as the lung remain poorly understood. Most studies characterizing the impact of aging on lung immunity have been conducted in mouse models or using clinical samples procured from individuals with chronic diseases (24). Data from these studies indicate functional deficits in alveolar macrophages (AM) including a refractory response to IFN*γ*, decreased phagocytosis, and increased basal cytokine production (23, 25). Additionally, these studies have reported the presence of dysfunctional CD8+ tissue resident T-cells (Trm) that are unable to adequately respond to viral challenges (26). However, data from rodent studies are confounded by their specific pathogen free (SPF) status. Unlike SPF rodents, humans are exposed to respiratory pathogens and irritants throughout their life, which shape the microbial and immunological landscapes of the lung. Clinical studies have reported an increased proportion of neutrophils and both reduced and increased abundance of macrophages in bronchoalveolar lavage (BAL) in aged individuals compared to younger adults (27, 28).

Over the past 10 years it has become clear that the lung harbors a diverse microbial community which is primarily composed of oral microbes such as *Prevotella, Fusobacterium, Neisseria, Streptococcus* and *Corynebacterium* (29, 30). However, the metabolic activity and ability of this community to self-renew remains contentious (31, 32). The composition of lung microbiome has been associated with environmental exposures (33), changes in pulmonary function (34), and shifts in local immunity (35). Changes in the lung microbiome have also been reported with multiple pulmonary disease states including asthma, idiopathic pulmonary fibrosis and chronic obstructive pulmonary disease (COPD) (36–39). For example, asthma is broadly associated with an increased abundance of Proteobacteria in the lung, driven by increased abundance of *Haemophilius*, *Neisseria,* and occasionally other pathogens such as *Pseudomonas* and *Klebsiella* (40–43). Similar patterns have also been observed in the lung microbiome of COPD patients experiencing an exacerbation which has decreased diversity and increased Proteobacteria driven by *Haemophilius* and *Acinetobacter* (38, 44). Despite dramatic increases in many microbiome-associated pulmonary disorders with advanced age, shift in the lung microbiome with healthy aging alone remains poorly defined. A recent study that used the sputum microbiome as a proxy for the lower-respiratory tract reported increased Firmicutes and a corresponding decrease in Proteobacteria with age (34). Additional studies have shown a loss of compartmentalization of upper respiratory tract (URT) communities with advanced age (45–47).

The mechanisms underlying increased susceptibility to respiratory infections are multifactorial and complex. Uncovering these mechanisms using only clinical specimens is challenging due to the difficulties associated with collecting respiratory specimens such as bronchoalveolar lavage samples from healthy humans and the challenges of disentangling the contribution of other confounding factors such as diet and smoking. Non-human primates (NHP) are an excellent animal model to carry out these types of studies for the following reasons. First, NHP share significant genetic, physiological similarities to humans and are genetically diverse. Second, outdoor-housed rhesus macaques such as the animals included in this study experience a life-long exposure to environmental and microbial antigens. Third, aged rhesus macaques recapitulate the physiological and systemic immunological hallmarks of immune senescence including the depletion of circulating naive T-cells, accumulation of memory T-cells, lymphoid fibrosis and, chronic low-grade inflammation (48–52). Fourth, rhesus macaques have been used to study microbiome development and stability at sites difficult to access in humans such as throughout the GI tract and within the lung (53–56). Fifth, rhesus macaques are already considered to be the gold standard model for the study of multiple human respiratory pathogens such as tuberculosis, high pathogenic influenza, and SARS-CoV-2 (57–59).

Therefore, in this study we carried out a comprehensive analysis of the impact of age on the immunological landscape, microbial community, and pulmonary function using adults (4.2 to 9.9 years old) and aged (18 to 29.0 years old) rhesus macaques. We report that the microbiome of aged animals becomes more heterogeneous across multiple sites. Additionally, as previously reported, a *Tropheryma-*dominated lung microbiome is common (∼50%) in rhesus macaques, especially adult animals. Since *Tropheryma* is known to infect alveolar macrophages, we stratified our immunological analysis by both host age and *Tropheryma* status. Aging led to an increase in the frequency of myeloid cell populations at the expense of T-cells. Transcriptionally we found that aging led to a shift from *GRZMB*+ to *IFNG*+ CD8+ T-cells. We also observed a decrease in *IL1B* transcripts in AM with age while expression of the senescence gene *FN1* increases. Together our findings lay the groundwork for understanding the impact of healthy aging on the lung and provide clues to increased respiratory infection severity with age.

## RESULTS

### Rhesus macaques display the circulating hallmarks of immunological aging

We first profiled circulating immune cell populations for known hallmarks of aging. Frequencies of total CD4+ and CD8+ T-cells were comparable between age-groups, while those of total B-cells and NK cells were decreased, and those of dendritic cells (DCs) and monocytes increased with age (Supp. **Figure 1A,B**). As expected, significant loss of naive CD4+ and CD8+ T-cells that was accompanied by an increase in both the central (CM) and effector memory (EM) cell in CD4+ T-cells and EM in CD8+ T-cells was observed with age (**Supp. Figure 1C,D**). Interestingly, frequency of naive and memory B cells was increased while that of marginal zone B-cells decreased with age in this cohort (**Supp. Figure 1E**). No differences in the frequency of DC cell subsets, non-classical CD16+ monocytes, or CD16+ NK cells were observed with age (**Supp. Figure 1F-H**). We also profiled changes in functional responses with age. Frequency of TNFα-producing monocytes was significantly increased with age in response to stimulation with bacterial and viral toll-like receptor (TLR) agonists (**Supp. Figure 1I**). No differences were observed in IFN*γ*/TNF*α* production by CD4+ and CD8+ T-cells following 16hr PMA stimulation. However, cytokine production by T-cells following PBMC stimulation with viral and bacterial lysate was increased with age (**Supp. Figure 1J,K**). Together, these data indicate that our study population exhibited canonical hallmarks of immune senescence and inflammaging (18–20).

**Figure 1:**
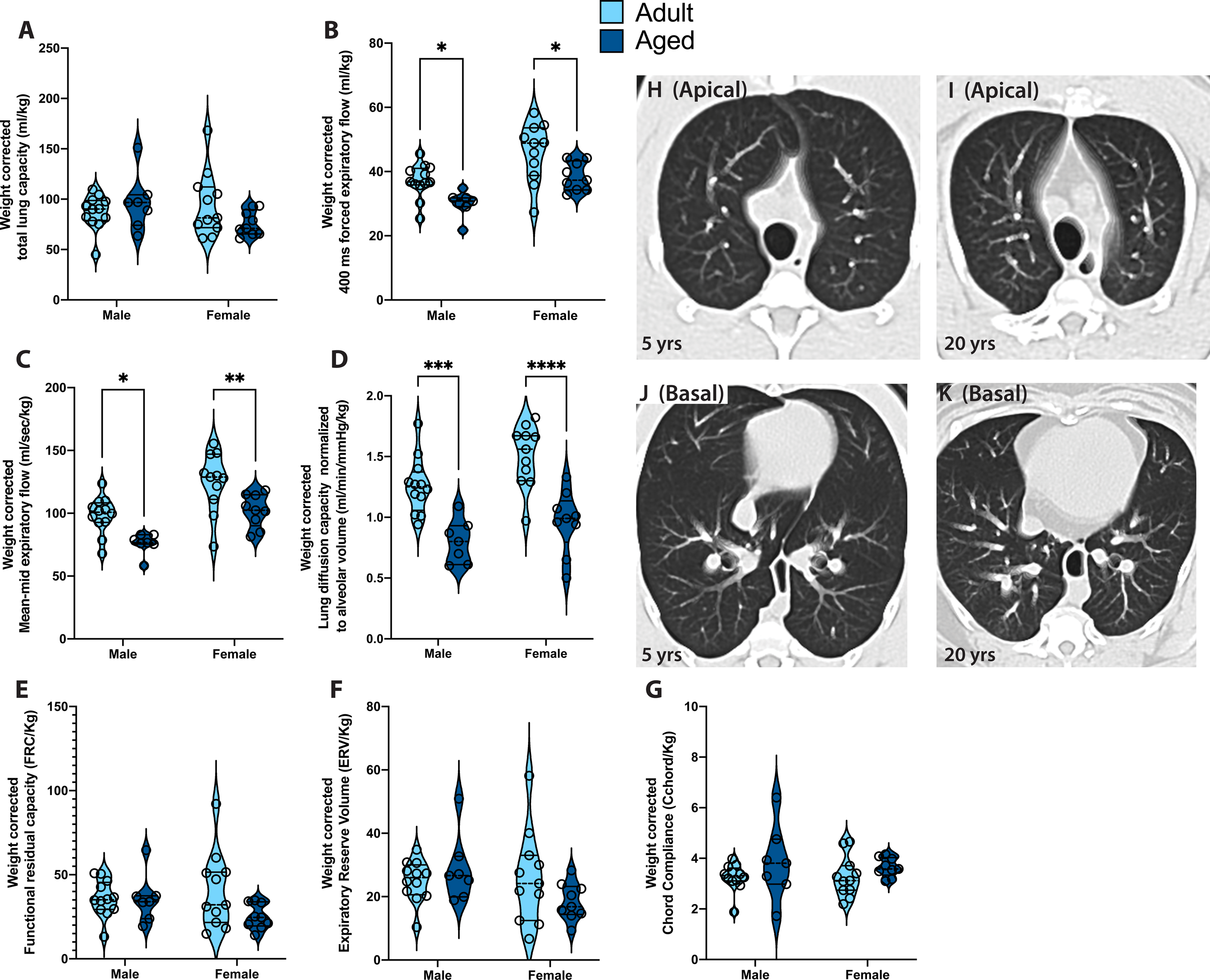
Age and sex related changes in pulmonary function. Pulmonary function tests were carried out in adult and aged rhesus monkeys and measures were normalized to animal weight. (A) Total lung capacity (TLC); (B) Forced expiratory flow in first 400 msec (FEV400) equivalent to the clinical FEV1 measure; (C) Mean-mid expiratory flow (MMEF, average expiratory flow rate between 25 and 75% of forced vital capacity); (D) Lung diffusion capacity normalized to alveolar volume (DLCO/VA); (E) Functional residual capacity (FRC); (F) Expiratory reserve volume (ERV); (G) Chord compliance (Cchord). Impact of age and sex and the interaction of these two factors on pulmonary function was determined by two-way ANOVA with the addition of Sidak post-hoc tests to determine the impact of age within each sex. Full ANOVA results can be found in **Supplemental Table 2** and significant (p < 0.05) post-hoc comparisons are displayed in each panel (* = p<0.05, ** = p<0.01, *** =p<0.001, **** =p<0.0001). (H-K) Apical and basal 5-slice projections (from 64 slice scans) from representative breath-hold CT scans from a 20 year and 5 year old rhesus monkey. All images were read by a thoracic radiologist who obvserved no differences in fibrotic or airway emphysematous changes between age groups.

### Aging leads to reduced lung function

Since respiratory disease results from the interaction of immune function with lung function, we next examined the effects of aging on lung function. A complete battery of lung function tests was performed to look at lung volumes, resistance and compliance, forced expiratory flows and volumes and gas exchange as we have previously described in other rhesus macaque studies (60, 61). Most measurements of lung function are highly related to height (17, 62), and for rhesus, weight is typically used as a surrogate for height, so all data is presented as absolute values and normalized for weight (**Figure 1, Supp. Table 2**). Total lung capacity (TLC) increased with age in males, which had significantly greater lung capacity than females (**Supp. Table 2**); however, when corrected for weight, no significant differences were noted with age or sex (**Figure 1A**). Forced expiratory flows, mean-mid expiratory flow and gas exchange capacity was comparable between adult and aged animals regardless of sex (**Supp. Table 2**), but when corrected for weight, forced expiratory flows (**Figure 1B,C**) and gas exchange (**Figure 1D**) were all significantly decreased with age in both males and females. No changes with aging were observed for Functional Residual Capacity (FRC), Expiratory Reserve Volume (ERV) or measures of static compliance (**Figure 1E-G).** Thoracic CT scans with controlled lung inflation in a subset of animals showed no emphysematous or fibrotic differences between adult (n=5) versus old (n=6) animals (**Figure 1 H-K**). These findings accurately recapitulate changes in human pulmonary function supporting the use of our model, and the combination of decreased pulmonary function and decreased gas exchange could work together to increase risk of respiratory infection in the aged.

### The lung microbiome of Rhesus macaques is often dominated by a single microbe

To determine the impact of aging on the lung bacterial community, we profiled the bronchoalveolar lavage (BAL) microbiome using 16S rRNA gene amplicon sequencing. In total 16 Amplicon Sequencing Variants (ASVs) were found in extraction (DNA extracted from PBS) and PCR negative controls and samples (**Supp. Figure 2A).** Of these, many ASVs were environmental contaminants (higher in negative controls than in samples) and were most commonly assigned to Comamonadaceae, Stenotrophomonas, Roseobacter and Alcaligenaceae; all of which were found in low abundance within BAL samples **(Supp. Figure 2A).** Additionally, we observed some ASVs in negative controls that were highly prevalent in our biological samples that are likely due to cross-ccontamination between PCR reactions from biological samples to negative controls such as Streptococcus ASV_535, Prevotella ASV_912, and Faecalibacterium ASV_950 (**Supp. Figure 2A).** These “cross-contaminates” were far more abundant in biological samples than negative controls. Overall, the contribution of environmental taxa to the composition of the BAL microbiome was minimal.

**Figure 2:**
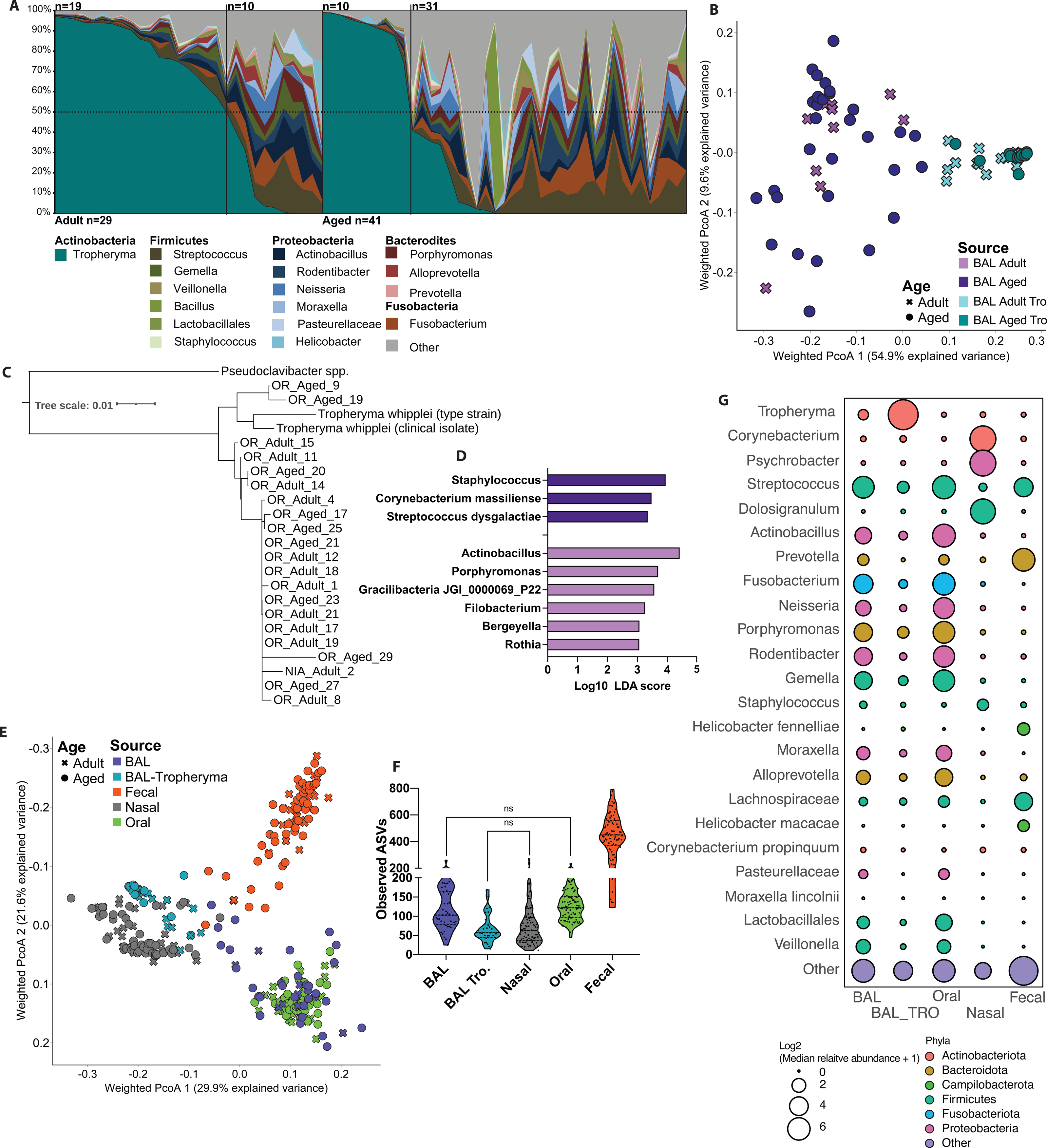
*Tropheryma* colonization and composition of the rhesus macaque lung microbiome. (A) Moving average of lung microbiome composition across all animals at the genus level. Any taxa found at <1% average abundance across samples grouped into the “other” category. (B) Principal coordinate analysis of weighted UniFrac distance from BAL samples, colored by host age, and *Tropheryma* dominated/non-dominated community types. (C) *Tropheryma* phylogenetic tree generated from the alignment of full length 16S rRNA gene sequences. (D) Differentially abundant bacterial taxa between age groups. Differential abundance was determined using LEFsE (Log10 LDA score > 2) of L7 (species level) taxonomy. € Principal coordinate analysis of weighted UniFrac distance generated from 16S amplicon microbiome data, colored by sampling site and shaped based on host age. (F) Violin plots of observed amplicon sequencing variants (ASVs) across sampling sights. All comparisons not denoted as NS on the figure were significant (p<0.05). (G) Bubble plot of the most abundant genera of bacteria across sampling sites. The size of each bubble represents the mean relative abundance for that site and color denotes which Phyla each genus is a member of.

Next, we probed the taxonomic landscape of the lung microbiome to determine the most abundant community members. Surprisingly, we found that a single genus (*Tropheryma*) dominated a large subset of samples making up >50% of the overall community in ∼2/3 of adult animals and ∼1/4 of aged animals (**Figure 2A**). In animals without *Tropheryma*-dominated community we observed a high relative abundance of the genera *Streptococcus*, *Fusobacterium*, and *Actinobacillus* among the 17 genera that were found at > 1% average abundance across all samples (**Figure 2A**). Interrogation of overall community composition using principal coordinate analysis of weighted UniFrac dissimilarity showed greater intra-group variability in non-*Tropheryma* dominated communities from aged animals compared to their adult counterparts (**Figure 2B, Supp Figure 2C**). To better classify the *Tropheryma*, we sequenced the full-length 16S rRNA gene from a subset of samples and compared it to the cultured *Tropheryma whipplei* type strain and other clinical isolates. The majority of *Tropheryma sp.* sequenced from the rhesus macaques BAL samples were highly similar and on average ∼97% similar to the *T. whipplei* type strain (**Figure 2C**). Next, we determined which bacterial taxa were differentially abundant between adult and aged animals without a *Tropheryma*-dominated community. Only species (Level 7 taxonomy) detected in at least 10% of samples at greater than 0.1% relative abundance within the sampled site were included in this analysis. No differentially abundant taxa were observed in *Tropheryma* dominated animals; however, several differentially abundant species were detected in non-*Tropheryma* dominated BAL communities (**Figure 2D**). Specifically, aging led to increased abundance of *Staphylococcus, Streptococcus dysgalactiae* and *Corynebacterium massiliense*, and reduced abundance of *Actinobacillus, Porphyromonas*, and *Rothia* in non-*Tropheryma* dominated BAL samples (**Figure 2D**). Of note, there was no correlation of pulmonary function with *Tropheryma* colonization.

The lung microbiome in humans is believed to be primarily composed of bacteria descending from the upper respiratory tract (URT) and more specifically the oral cavity (29, 30). Moreover, studies have shown a loss of tight compartmentalization with age in the URT (45–47). Therefore, we sequenced paired nasal, oral, and fecal samples from all animals. We initially analyzed this dataset agnostic of host age to profile microbiome biogeography. The overall composition of the microbiome at these three sites were dissimilar due to a combination of distinct bacterial genera and difference in alpha diversity at each site (**Figure 2E-G**). As expected, the number of amplicon sequencing variants (ASVs) was significantly higher in fecal samples compared to the BAL, nasal, and oral communities (**Supp Figure 2B**). The oral bacterial community was dominated by *Streptococcus* and *Fusobacterium* while, the nasal microbiome was dominated by *Corynebacterium*, *Psychrobacter,* and *Dolosigranulum* that were largely absent at all other sites (**Figure 2G**). Finally, fecal samples had a high abundance of *Prevotella*, *Helicobacter*, and *Streptococcus* (**Figure 2G**).

We next determined how the microbial community of the lung related to other sampled sites. The *Tropheryma*-dominated lung microbiomes were phylogenetically most similar to nasal microbiomes due to high abundance of Actinobacteria (albeit different species) and low alpha diversity rather than shared bacterial genera (**Figure 2E, Supp Figure 2B**). Conversely the non-*Tropheryma* lung and oral microbiomes shared several genera (*Streptococcus*, *Actinobacillus*, *Fusobacterium*) (**Figure 2E**). We next examined which ASVs were shared and unique between lungs and other body sites (**Supp Figure 3 A-C)**. Only ASVs detected within at least 10% of samples at >0.1% relative abundance at the indicated sites were included in this analysis. Surprisingly, a large number of ASVs were found to be unique to both *Tropheryma*-dominated and non-*Tropheryma* BAL microbiomes on average accounting for 16.5 and 82% of the total community respectively **(Supp Figure 3A-C)**. As expected, *Tropheryma* (ASV_4722, ASV_4310) was the most abundant lung-specific microbe **(Supp Figure 3B,C)**. Additional abundant lung-specific microbes included *Porphyomonas canoris* ASV_1309, *Prevotella* ASV_2044, *Acinetobacter* ASV_4580, and Comamonadaceae ASV_3409 **(Supp Figure 3B,C)**. A large portion of ASV’s identified in BAL samples were also identified in the oral cavity (**Supp Figure 3A**). However, the abundance of these shared microbes varied widely between *Tropheryma*-dominated and non-*Tropheryma* animals (**Supp Figure 2B-D)**. A large portion of the BAL microbiome of non-*Tropheryma* animals was composed of ASVs shared primarily with the oral microbiome notably, *Gemella* ASV_3506, *Porphyromonas* ASV_761 and *Fusobacterium* ASV_1429 (**Supp Figure 3A-C**). In contrast, a small number of ASVs were shared between the BAL and nasal or fecal microbiomes and these ASVs were found at low abundance (**Supp Figure 3A-C)**. Additionally, some ASVs were shared between the BAL and oral microbiomes as well as at least one other site such as *Neisseria* ASV_4242 and *Moraxella* ASV_338 (BAL, oral nasal) along with *Actinobacillus* ASV_1196 and Streptococcus ASV_535 (BAL, oral, fecal) **(Supp Figure 3A)**. Finally, three ASVs were shared across all sites, with *Streptococcus* ASV_4404 being the most abundant **(Supp Figure 3A-C)**. Together these data suggest that while a large percentage of the BAL microbiome is shared with the oral cavity, it also harbors a significant unique microbial community.

**Figure 3:**
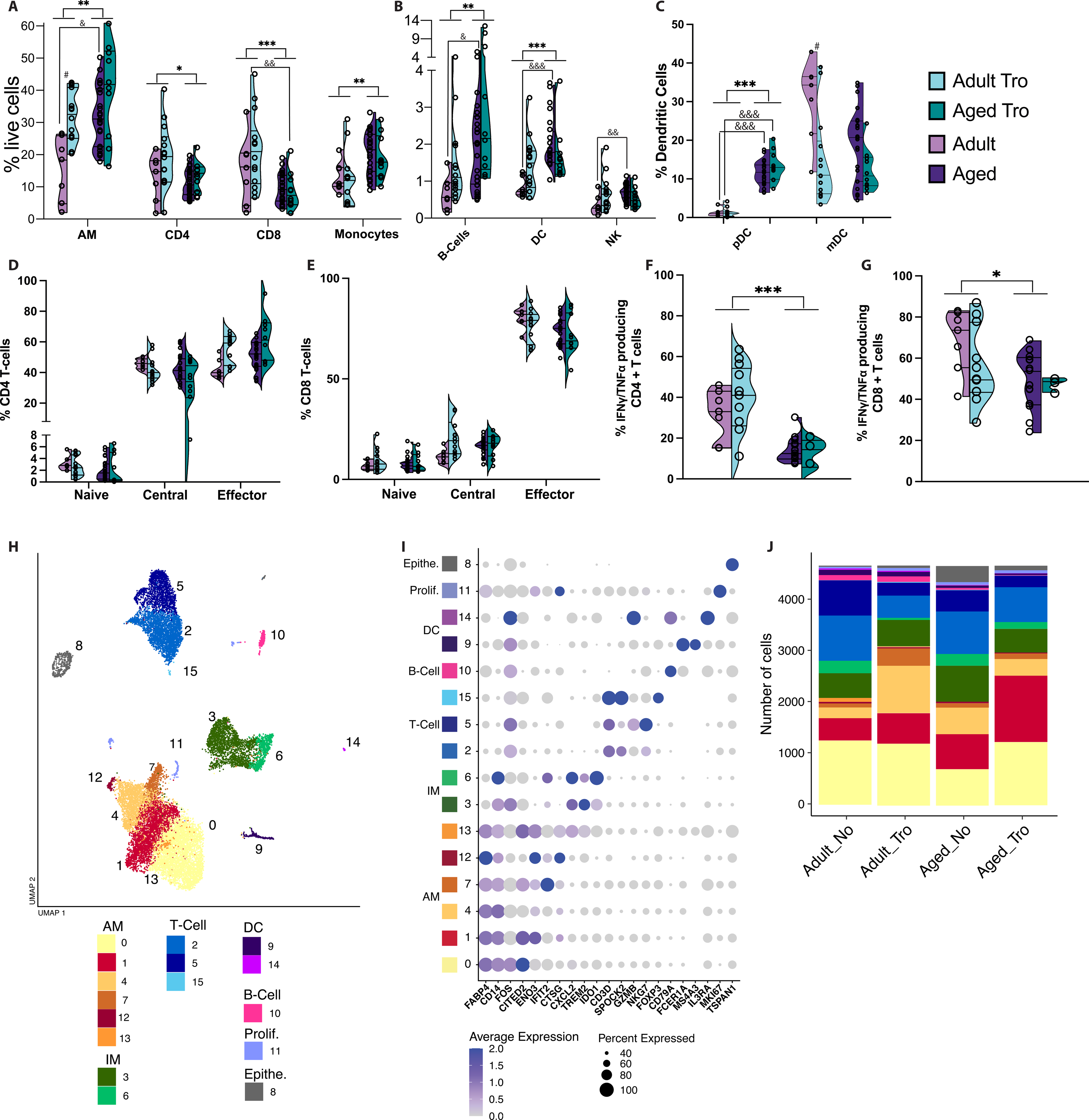
Age-related differences in lung immune cell frequency and function. Violin plots of immune cell frequency measured by flow cytometry of BAL cells for (A) major immune cell subsets, (B) CD16+ Monocytes, (C) Plasmacytoid and Myeloid Dendritic cells, (D) CD4+ T-cell subsets, a€(E) CD8+ T-cell subsets. (F,G) Percent abundance of IFN*γ* and/or TNF*α* producing CD4+ and CD8+ T-cells after 16hr PMAi stimulation. Cytokine producing cells were measured using intra-cellular cytokine staining and flow cytometry. Comparisons were made using an unpaired t-test between age-groups for each cell subset independent of *Tropheryma* (Tro.) colonization (* = p<0.05, ** = p<0.01, *** =p<0.001, **** =p<0.0001). Additional comparisons were made between Adult vs. Adult Tro. (# = p<0.05) , Aged vs. Aged Tro. (# = p<0.05), Adult vs. Aged (& = p<0.05), and Adult Tro. Vs. Aged Tro. (& = p<0.05). (H) UMAP projection of all 18,712 BAL cells. Colored by cluster with major immune cell subsets are annotated. (I) Bubble plot of identifying genes highly expressed in each cluster determined using the FindMarkers function in Seurat. Level of normalized expression is shown using a color scale ranging from low (white) to high (purple). Size of the bubble is representative of the fraction of cells within each cluster expressing the marker. (J) Stacked bar plot depicting the absolute abundance of cell clusters withing each group. AM = alveolar macrophage, IM = infiltrating macrophage, DC = dendritic cell, Prolif. = Proliferating cell, Epithe. = epithelial cell.

Next, we determined the impact of aging on microbial communities within all three sites distal to the lung. We found that aging did not lead to changes in observed amplicon sequencing variants (ASVs), a measure of microbiome richness at any site (**Supp Figure 2B).** However, aging was associated with a higher within-site dissimilarity in the nasal, fecal and non-*Tropheryma* BAL samples and a reduction with age in *Tropheryma* BAL (**Supp. Figure 2C**). Next, we determined which bacterial taxa were differentially abundant between adult and aged animals at each site. Only species (Level 7 taxonomy) detected in at least 10% of samples at greater than 0.1% relative abundance within the sampled site were included in this analysis (**Supp. Figure 2D-F**). We observed an increase in pathobiont Proteobacteria such as *Escherichia-Shigella* and *Campylobacter corcagiensis* in the fecal samples of aged animals, while abundance of *Lactobacillus* and *Treponema* was decreased (**Supp. Figure 2D, Supp. Table 3**). Interestingly some species were differentially abundant in aged animals across multiple sites including *Streptococcus dysgalactiae, Corynebacterium massiliense, and Corynebacterium propinquum* (**Figure 2D, Supp. Figure 2D-F**).

### Impact of aging on the pulmonary immunological landscape

Next, we assessed the impact of aging and *Tropheryma*-dominant microbiome on immune cell populations in the lungs. Analysis of major populations in the BAL revealed significant differences in frequencies of alveolar macrophages (AM), infiltrating monocytes/macrophages (IM), T cells, B cells, NK cells, and DCs with both age and *Tropheryma* status (**Figure 3 A,B**). Specifically, frequencies of AM, IM, DC and B-cells increased while those of CD4+ and CD8+ T-cells decreased with age (**Figure 3 A,B**). Many of these patterns were maintained when the age groups were also split by *Tropheryma* status (**Figure 3 A,B**). The increased frequency of DCs was mediated by an increase of plasmacytoid dendritic cells (pDC) in aged animals, while the presence of *Tropheryma*-dominated community was associated with decreased frequency of mDC in adult animals (**Figure 3C**). No differences were observed in the breakdown of CD4+ or CD8+ T-cells memory subsets (**Figure 3D,E**). To determine the impact of aging on functional responses, BAL cells were stimulated with PMAi for 16 h, and cytokine production was measured by intracellular cytokine staining. The frequency of INF*γ*/TNF*α* producing CD4+ and CD8+ T-cells was reduced in aged animals regardless of *Tropheryma* status (**Figure 3F,G**). In summary, aging results in a remodeling of the immunological landscape with a reduction in both the frequency and functional capacity of T cells while myeloid cells became dominant.

To gain a deeper understanding of the impact of aging on the immunological landscape, BAL cells from 12 animals (n=3/group with each group pooled into a single sequencing library) were profiled using scRNA-Seq. Although this pooling strategy ensured biological diversity, the lack of hashing precluded us from conducted statistical analyses of changes in subset distribution. Dead cells and doublets were removed then each group was down-sampled to 4678 cells prior to downstream analysis to compare absolute abundance of cell clusters rather than relative abundance. Principal component analysis and dimensional reduction using Uniform Manifold Approximation and Projection (UMAP) revealed distinct clustering into major myeloid and lymphocyte subsets (**Figure 3H**). Clusters were annotated based on highly expressed marker genes using Seurat’s FindMarkers function (**Figure 3I and Supp. Table 4**) This approach revealed changes in proportions of AM, IM, as well as major T and B cell subsets with age and *Tropheryma* status, as well as shifts in AM cell states (**Figure 3J**).

### Aging leads to functional shifts in BAL T-cells

T cell subsets were re-clustered to identify changes in cell states with higher resolution (**Figure 4A**). This led to the identification of 7 clusters that can be partitioned into CD8+ T-cells (clusters 1 and 3), CD4+ T-cells (clusters 0 and 5), Gamma-delta T-cells (*γδ* T-cells; cluster 2), regulatory T-cells (Tregs; cluster 7), and NK cells (cluster 6) based on the expression of established marker genes (**Figure 4A,D**). Both aging and *Tropheryma* colonization led to a reduction of total T/NK cells (**Figure 4B**). In the absence of *Tropheryma* colonization, this reduction was driven by a loss of *ITK/IL7R* CD4 T-cells (cluster 5) (**Figure 4B,D)**, and expansion of *γδ* T-cells (cluster 2), and CD8 T-cells (clusters 1 and 3) independent of age (**Figure 4B,D)**.

**Figure 4:**
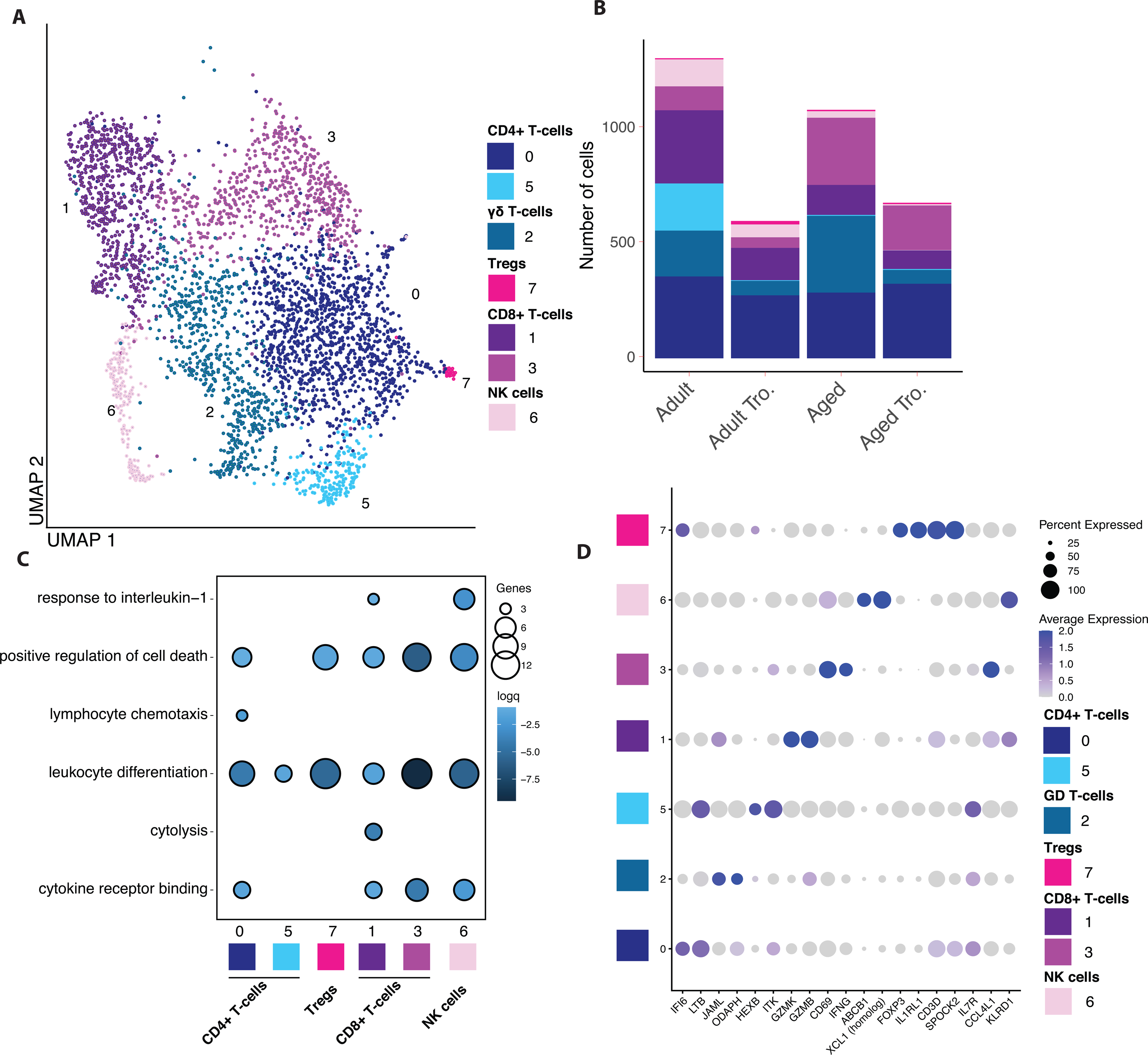
Aging leads to a reduction of GZMK/GZMB expressing and increase of IFNG/CCL4L1 expressing CD8+ T cells. (A) UMAP projection of T and NK cells re-clustered from the main UMAP. Major cell types are labeled. (B) Stacked bar plot depicting the absolute abundance of T and NK cell clusters withing each group. (C) Functional enrichment of genes differentially expressed each T and NK cell cluster when compared to all other clusters. Color and size of the bubble represents statistical significance and number of genes respectively. Differentially expressed genes determined using the Seurat FindMarkers function. (D) Bubble plot identifying genes highly expressed in AM and IM cell clusters determined using the FindMarkers function in Seurat. Level of normalized expression is shown using a color scale ranging from low (white) to high (purple). Size of the bubble is representative of the fraction of cells within each cluster expressing the marker.

We next carried out functional enrichment of the marker genes that defined each cell cluster, identified using the Seurat “FindMarker” command, to elucidate the functional potential of each cell cluster (**Figure 4C, Supp. Table 5**). As expected for tissue resident lymphocytes, marker genes that defined all CD8 T-cell clusters enriched to the GO term ”leukocyte differentiation” (e.g., *FOS, TNF, CD83*), and this enrichment was strongest for *IFNG* expressing CD8 T-cells (cluster 3) suggesting that this cluster is terminally differentiated (**Figure 4C**). Interestingly, all gene markers except for those defining cluster 5 (which was primarily found in non-*Tropheryma* adult animals) enriched to the GO term “positive regulation of cell death” (e.g., *CTLA4, CD40LG, NTRK1*) (**Figure 4C**). Genes that defined cluster 1 CD8 T-cells, which decreased with age, enriched to both “cytolysis” (e.g. *GZMH, GZMA, GZMB, GZMM*) and “response to interlukin-1” (e.g. *IL1B, CCL5, CCL4L1*), while markers of cluster 3 CD8 T-cells, which increased with age, enriched to “cytokine receptor binding” (e.g., *CSF1, IFNG, TNFSF9, IL21*) (**Figure 4C**), suggesting a potential shift in functional capacity of CD8 T-cells with age.

We next determined how aging impacts the transcriptional profiles of CD4, CD8 and *γδ*T-cells. To that end, cells within these subsets were concatenated and DEGs between adult and aged animals within each group (based on *Tropheryma* status) were identified as those with log2FC >0.25 and FDR p <0.05 (**Supp Figure 4A-E**). Within BAL samples obtained from animals without a *Tropheryma*-dominated community, CD4 T cell DEGs enriched predominantly to GO terms associated with “chemotaxis” (e.g., *CCR6, CCL4L1*) and “apoptosis” (e.g., *GZMB*) while *γδ* T-cells DEGs enriched predominantly to GO terms associated with “migration” (e.g. *CCL5*) and “cytokine signaling” (e.g. *IFNG, LTB*). CD4 T cell DEGs from *Tropheryma*-dominated BAL samples enriched to “activation” (e.g. *CXCL13, CD40LG*), “apoptosis” (e.g. *CTSD, EGLN3*), “response to bacterium” (e.g. *LYZ, IL1B*), and “type-I interferon signaling pathway” (e.g. *IRF1, ISG15*) (**Supp Figure 4A,B**). *γδ* T-cells DEGs from *Tropheryma*-dominated BAL samples enriched to GO terms associated with “metabolism” (e.g. *SLC2A1, SLC2A3, NR4A3*),” cell adhesion” and “signaling” (e.g. *TMIGD2, ITGA6*) (**Supp Figure 4A,C**).

Within CD8 T cells, DEGs enriched predominantly to GO terms “immune activation”, “antiviral immunity”, and “apoptosis” (**Supp Figure 4D**). Overall, genes associated with immune activation were significantly upregulated with age regardless of *Tropheryma* status, notably *IFNG*, *TNF* and *NFKBID* (**Supp Figure 4E**). Genes important for the response to oxidative stress such as *SOD2* and *ID3* were downregulated with age (**Supp Figure 4E**). Together these data suggest that independent of *Tropheryma* colonization, aging leads to large shifts in BAL CD8 T-cell functional potential, while *Tropheryma* colonization is associated with a bigger impact on gene expression within CD4 and *γδ* T-cells.

### Aging and *Tropheryma* status impact the transcriptional profile of Macrophages

In both humans and rhesus macaque, AM and IM are the most abundant immune cells in the lung. To explore how aging and *Tropheryma* status is associated with this critical population, we re-clustered the myeloid cells (**Figure 5A,B**). This analysis revealed 6 cell states for AM (clusters 0, 1, 3, 4, 5 and 7 expressing high levels of *FABP4* and *MARCO*) (**Figure 5A, Supp Figure 5A**). Within the AM subsets, clusters 0, 1 and 4 express markers consistent with a self-renewing regulatory population (*RGCC*, *SOCS1* and *ENPP1*) (**Supp Figure 5A).** In contrast, AM cluster 7 expressed high levels of interferon stimulated genes *IFIT2* and *MX1* suggesting this subset may play a role in anti-viral defense (**Supp Figure 5A**). *Tropheryma* colonization led to increased abundance of AM, especially clusters 3 and 4 in adult animals and cluster 5 in aged animals (**Figure 5B**). Additionally, we observed age- and *Tropheryma*-dependent transcriptional patterns across clusters. For example, expression of *IL1B*, a key inflammatory mediator, was decreased with age across all clusters while that of the pro-fibrotic gene *FN1* was increased (**Figure 5D**). Moreover, expression of the complement factor *C1QC* was higher in AM from *Tropheryma*-colonized animals across clusters (**Figure 5D**). Finally, expression of *CD69*, a marker of early activation, was highest in adult animals that lack a *Tropheryma*-dominated community across all clusters, while *FOS*, a suppressor of the inflammatory response was up-regulated with age and *Tropheryma* status (**Figure 5D**). Within IM, cluster 2 was most abundant across all groups and expressed high levels of the M2-like marker *TREM2,* complement gene *C1QC, and CPVL* (a gene that plays a critical role in MHC-I antigen presentation). On the other hand, cluster 6 expressed high levels of M1-like markers *IDO1* and *CCL3* (**Figure 5B,E, Supp Figure 5A**). Abundance of cells in cluster 6 was reduced with aging and *Tropheryma* colonization (**Figure 5B**) while *FCGR3* (CD16) was upregulated in both *Tropheryma* colonized groups in both IM clusters (**Figure 5E**).

**Figure 5:**
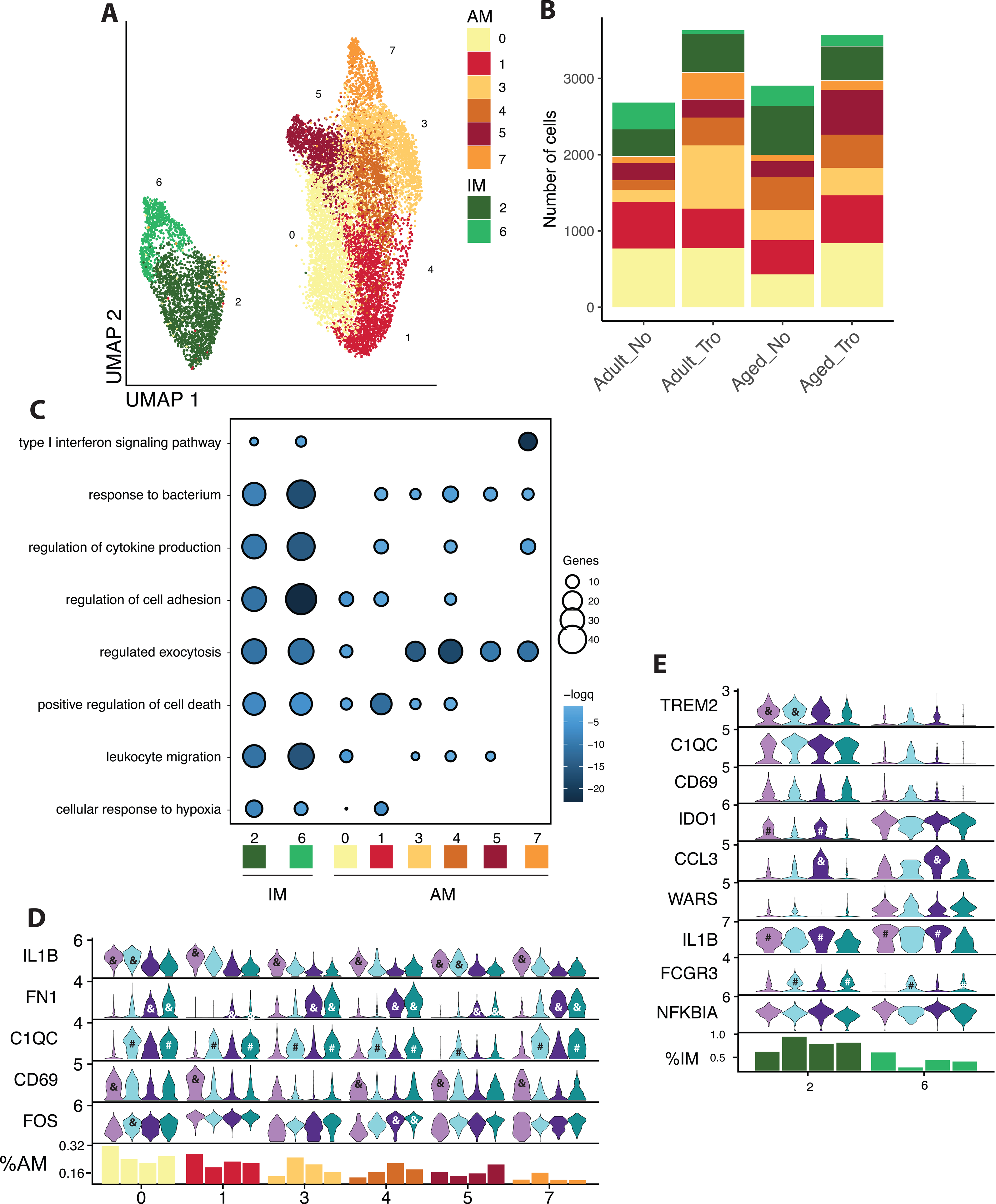
Aging leads to increased fibronectin 1 expression in AM and shift in expression of IL1B from AM to IM. (A) UMAP projection of AM and IM cells re-clustered from the main UMAP. Major cell types are labeled. (B) Stacked bar plot depicting the absolute abundance of AM and IM cell clusters withing each group. (C) Functional enrichment of genes differentially expressed each AM and IM cell cluster when compared to all other clusters. Size of the bubble represents statistical significance and number of genes respectively. Differentially expressed genes determine using the Seurat FindMarkers function. (D,E) Violin plots depicting the expression of key genes between groups across AM (D) and IM (E) cell clusters. Violins colored by group. Bar graph at the bottom of the figure represents the relative abundance of each cluster with the indicated group as a percentage of the total AM (D) and IM (E) cells. For D and E significance was determined for each gene within each cell clusters using the non-parametric Kruskal-Wallis ANOVA with Dunn’s multiple comparison test for post-hoc pairwise comparison when the initial ANOVA was significant. Post-hoc significance between Adult vs. Adult Tro. (# = p<0.05) , Aged vs. Aged Tro. (# = p<0.05), Adult vs. Aged (& = p<0.05), and Adult Tro. Vs. Aged Tro. (& = p<0.05) are show in each panel.

Functional enrichment was then carried out using marker genes that defined each cluster to infer differences in functional potential between the various cell states (**Figure 5C, Supp. Table 6**). Marker genes enriched to similar GO terms; however, genes that define IM enriched more significantly to most GO terms (e.g., “response to bacterium”, “regulation of cytokine production”, “leukocyte migration”) (**Figure 5C**). Marker genes of AM Cluster 1 enriched more significantly to GO term “positive regulation of cell death” while clusters 3-7 enriched more significantly to “regulated exocytosis”, and cluster 7 genes to “type 1 interferon signaling pathway” (**Figure 5C)**.

We next investigated the impact of aging and *Tropheryma* colonization on transcriptional profile of AM and IM. Overall, aging in the absence of *Tropheryma* colonization resulted in a more profound transcriptional shift (**Supp Figure 5B**). Within infiltrating macrophages, DEGs enriched to GO terms such as “leukocyte activation involved in immune response” and “response to bacterium” (**Supp Figure 5B**). Specifically, inflammatory genes such as *TNF, S100A8, IL1B, RGS1 and NFKBIA* were significantly upregulated with age, while genes that encode anti-inflammatory proteins (*CHIT1*, *FTL*) and play a role in endocytosis (*CTSB*, *CTSD*, *SPP1*) were downregulated (**Supp Figure 5C**). Interestingly, expression of *CXCL1*, *CD63*, and *CCL17* within infiltrating macrophages showed opposite but significant age-related expression changes based on the *Tropheryma* status (**Supp Figure 5C**).

Genes that are differentially expressed with aging in AM enriched to GO terms such as “leukocyte degranulation” and “response to bacterium” (**Supp Figure 5B**). At the DEG level, multiple pro-phagocytic and inflammatory genes (e.g. *IL1B, CXCL1, CCL4L1, IDO1, CCL2 and SOD2*) were downregulated, while genes important for tissue homeostasis (e.g. *LGALS3, GRN, LCN2, MRC1, GPNMB*) were upregulated with aging (**Supp Figure 5D**). In AM from animals with a *Tropheryma*-dominated BAL bacterial community, genes associated with apoptotic signaling (e.g., *BAG3*, *ATF3*) were upregulated, coupled with a downregulation of interferon stimulated genes (e.g., *IFI27*, *IGHM*, *ISG15*) (**Supp Figure 5D**). Finally, expression of *S100A8*, and *S100A9*, which promote inflammation, was increased across all BAL macrophages with age (**Supp Figure 5C,D**). Together these data suggest that aging leads to the upregulation of inflammatory and pro-fibrotic genes, and down-regulation of genes important for tissue homeostasis and antigen uptake and processing. These changes could lead to reduced anti-microbial defense but heightened aberrant inflammation.

## DISCUSSION

While systemic markers of immunosenescence and inflammaging have been well explored, there is a paucity of studies exploring patterns of immunological aging in tissues, particularly the lungs. In this study we sought to determine the impact of healthy aging on the functional, microbial, and immunological landscapes of the lung using the rhesus macaque model. Our understanding of age-mediated changes in the lung immunological, functional, and microbial landscape remains limited due to the challenges associated with collecting BAL samples from healthy humans. Nonhuman primates, notably rhesus macaques, share significant genetics and physiological similarities with humans that make them a highly reliable pre-clinical model. Indeed, the rhesus macaque model recapitulates the hallmarks of systemic immune senescence described in humans (48, 52). We used BAL samples from adult and aged macaques and a multi-pronged approach to interrogate age-induced changes in lung immune status, microbial community composition and pulmonary function.

Our analysis of pulmonary function in aging rhesus demonstrates similar declines as seen in humans (12–17), including hallmark decreases in forced expiratory volumes, forced expiratory flows and gas exchange. These differences do not appear to derive from emphysematous changes or increased fibrosis as no such changes were seen in CT scans. This is not however definitive as no histologic analysis was performed and emphysematous changes have previously been reported in aged rhesus macaques (63). This decrease in respiratory capacity and alveolar function when combined with infection could increase the likelihood of clinically significant disease due to poor clearance of pathogens and warrants further investigation. Indeed, similar to humans, aged macaques suffer from more severe respiratory disease, including from SAR-CoV-2 and Influenza viruses (64, 65).The decrease in alveolar gas exchange capacity will similarly increase the likelihood of clinically significant decreases in blood oxygen levels with respiratory infection leading to increased tissue damage.

Interestingly, our analysis of the microbiome showed that several animals had a *Tropheryma spp.* dominated BAL microbial community. Earlier studies that profiled the lung microbiome of macaques have also reported a high prevalence of *Tropheryma* (55, 56). Specifically, a longitudinal analysis showed that 50% of the animals had *Tropheryma*-dominated lung microbiome prior to SHIV infection. Moreover, following SHIV infection, 75% of animals had at least one *Tropheryma*-dominated sample during a 1-year follow up, suggesting that *Tropheryma* is a common member of the captive healthy macaque lung microbiome (56). In humans *Tropheryma* is most associated with Whipple’s disease, an infection in which *T. whipplei* infects macrophages of the small intestine (66, 67). However, *Tropheryma spp.* species can also be found in the human lung without any clear clinical symptoms, and their abundance in increased in HIV+ individuals (68, 69). Since the lung is one of the least sampled human tissues for microbiome composition, the prevalence of *Tropheryma* may be higher than currently estimated (70). Additionally, it is well established that gut and vaginal microbiomes of rhesus macaques are more similar to those of humans in the developing world than western countries (53, 71–73). It is possible, that *Tropheryma* lung colonization among humans could also vary in this manner.

Most animals in our study [41/70] had diverse lung microbial communities with little or no *Tropheryma* and were dominated by microbes typically found in the oral cavity. The oral cavity has been well established as the primary source of the lung microbiome in humans (30, 74). While it was initially hypothesized that the presence of oral microbes in the lungs was due to contamination during the bronchoscopy procedure, more in-depth studies have shown that this contamination is negligible and that oral microbes infiltrate the lungs primarily by micro-aspiration (29). As we and others have previously shown for the fecal, vaginal and oral microbiome (53, 71, 75), most microbes that colonize the lungs (i.e. *Streptococcus, Neisseria, Prevotella, Fusobacterium,* etc.) of rhesus macaques are also found in humans (29, 30, 74). One exception to pattern was the colonization of the rhesus macaque lung by *Rodentibacter*, which is commonly found in the oral microbiome of rodents but not humans (76).

Additionally, we show that aging leads to increased intragroup heterogeneity of microbial communities across multiple sites (Fecal, Nasal, non-*Tropheryma* BAL). This pattern has previously been observed in the gut microbiome of humans (77). The fact that this pattern is recapitulated in aged macaques in the absence of significant differences in diet and exposure to environmental pollutants and microbes strongly suggest other age-mediated changes are important (e.g., changes in immunological control and tissue elasticity). In agreement with human studies (46, 47, 78), we also observed an increased relative abundance of proteobacteria pathobionts (*Escherichia-Shigella* and *Campylobacter corcagiensis*) in the gut microbiome with aging. Moreover, multiple age-related differentially abundant bacterial taxa within the lungs of non-*Tropheryma* animals including *Streptococcus dysgalatiae, Corynebacterium massiliense, and Staphylococus* were identified in aged animals. Conversely, the lung microbiome of non-*Tropheryma* adult animals had an increased abundance of *Actinobacillus*, *Rothia*, and *Porphymonas* among others. Unexpectedly these taxa were not differentially abundant in the oral microbiome which is the most likely source of infiltration into the lung. This may suggest that while oral microbes are constantly infiltrating into the lungs, aging leads to an environment that is more permissive to colonization by specific microbes.

Alveolar macrophages (AM) are self-renewing and well-established as the primary orchestrators of the pulmonary immune system in both health and disease (79, 80). The loss or impaired function of AM has been shown to lead to increased pathology and mortality in mice following viral infection (81, 82). Despite their important role, little is known about the impact of healthy aging on these cells. Aging led to an increased abundance of AM in rhesus macaques. These data are in contrast to those reported in inbred mice (23) and a recent study in humans showing no changes in the abundance of airway macrophages (83). These discordant results may be due to the lack of life-long exposure to pathogens and environmental pollutants (in the case of SPF mice) and differences in sampling procedures in the case of clinical studies. Previous studies have found that pulmonary neutrophils increase with age and are a significant contributor to age related pulmonary inflammation (84); however, we were unable to assess changes in neutrophil frequencies as we used cryopreserved BAL samples.

We observed an increased expression of the complement factor *C1QC* across all AM clusters obtained from animals colonized by *Tropheryma*. Complement plays an important role in linking the innate and adaptive immune system in pulmonary disease (85), however, it is unclear whether *Tropheryma* infects AM and if complement is a mechanism by which this bacterium is controlled. Independent of *Tropheryma* colonization, we observed distinct shifts in the transcriptional pattern of AM with age. For example, AM from aged individuals had heightened expression of Fibronectin 1 (*FN1).* The expression of FN1 expression and other extracellular matrix proteins is tightly regulated in the lungs and under homeostatic conditions these protein play an important role in tissue repair (86). However, overexpression of these factors may lead to the development of idiopathic pulmonary fibrosis (87, 88) and acts as a predictor of resistance to chemotherapies in some cancers (89, 90). Interestingly, transcripts of *IL1B* were decreased within AM with age, which may indicate reduced immunosurveillance by AM in aged animals. In contrast to AM, expression of *IL1B, CCL3,* and *TNF* in IM increased with age in non-*Tropheryma* animals. The lack of TNF induction in IM from *Tropheryma* dominated samples is in line with reports of reduced TNF-expression by bystander macrophages in *T.whipplei* infected gut samples (91).

The increased abundance of AM and infiltrating macrophages was accompanied by a corresponding decrease in CD4+ and CD8+ T-cells in the BAL. Moreover, frequency of T-cells secreting cytokines in response to polyclonal stimulation decreased with age. From a transcriptional standpoint, CD8 T-cells in adult animals predominantly expressed high levels of Granzymes (*GZMK, GZMB*) and were predicted to carry out cytolytic functions. In contrast, CD8+ T-cells in aged animals expressed high levels of *IFNG* and their marker genes are predicted to play a role in differentiation and apoptosis. BAL CD8+ Trm are essential to control respiratory viral infection and can provide long-lived immunity (92, 93). However, dysfunctional CD8+ Trm response can also lead to damage during and after viral infection (26, 94). The reduced expression of granzymes genes in aged CD8 T-cells suggest reduced cytolytic capacity, a key factor for clearing viral pathogens (95, 96). On the other hand, IFN*γ* does not play a prominent role in CD8+ Trm mediated respiratory viral clearance (97–99) but may play a role in immunopathology (99). Indeed, previous studies of influenza in mice have found that viral infection causes a distinct accumulation of CD8+ Trm in aged animals (100). However, these cells lack key effector functions in mice (26). Our findings are consistent with a transcriptional shift with aging indicative of reduced anti-microbial function.

We observed the most pronounced age-related shift in the transcriptional profiles of BAL CD4+ T-cells in *Tropheryma* dominated animals, including the upregulation of genes associated with anti-bacterial responses (*LYZ, CXCL13*). Since *Tropheryma* colonization was not associated with clinical disease, these observations suggest that CD4 T cells may play a role in maintaining this homeostasis. Indeed, colonization of the lung by *Tropheryma* in humans is most common in HIV+ individuals who have reduced and dysfunctional CD4+ T-cells (68), and Whipple’s disease (an infection of the small intestine by *Tropheryma whipplei*) is associated with a reduced CD4+ Th1 response (101). CD4+ T-cells from the BAL non-*Tropheryma* animals showed limited age-related transcriptional changes. One notable exception is increased expression of *CCL4L1* in CD8+ and CD4+ T-cells. *CCL4L1* is subsequently translated into Macrophage inflammatory protein 1*β* (MIP-1*β*) a key chemoattractant of monocytes and essential for controlling pulmonary infections such as COVID-19 (102). However, in the absence of infection, increased T-cell production of MIP-1*β* increases with age may be associated with aberrant inflammation (103). Finally, we identified limited age-related transcriptional changes in BAL *γδ* T-cells; however, this cell population was depleted in both adult and aged *Tropheryma*-dominated animals. Given the cross-sectional nature of our study it is not possible to determine if this depletion is a cause or consequence of *Tropheryma* colonization.

Understanding how the lung environment changes with healthy aging is essential to combatting respiratory disease in a rapidly expanding aged population. In this study we compared BAL samples from adult and aged rhesus macaques to delineate the impact of “healthy aging” on immunological, microbial, and functional potential of the lung. Our key findings indicate that aging leads to a more inflammatory environment that may have compromised anti-microbial function. This change combined with the decrease in pulmonary function and gas exchange could increase the risk of clinically significant respiratory infections with aging. Independent of age, *Tropheryma* is a prevalent and dominant colonizer of the rhesus macaque lung and had a significant impact on pulmonary immune composition and function, despite the absence of respiratory symptoms. Given these findings, future studies should determine how differences in the lung microbiome may impact the host response to infection. Specifically in macaques, which serve as a gold standard model for multiple respiratory pathogens, the impact of *Tropheryma* colonization should be assessed. These data pave the way for studies aimed at designing and testing interventions to reduce or reverse age-associated shifts in the lung environment to reduce age-related pulmonary disease in the elderly.

## Acknowledgements

This work was supported, in part, by the intramural program of the National Institute on Aging, NIH. The authors thank the ONPRC Division of Comparative Medicine and, veterinary staff for their expert animal care. The authors also thank the veterinary technicians at the NIA nonhuman primate core for sample collection and support from the Division of Veterinary Resources animal care team.

---

**Figure S1:**
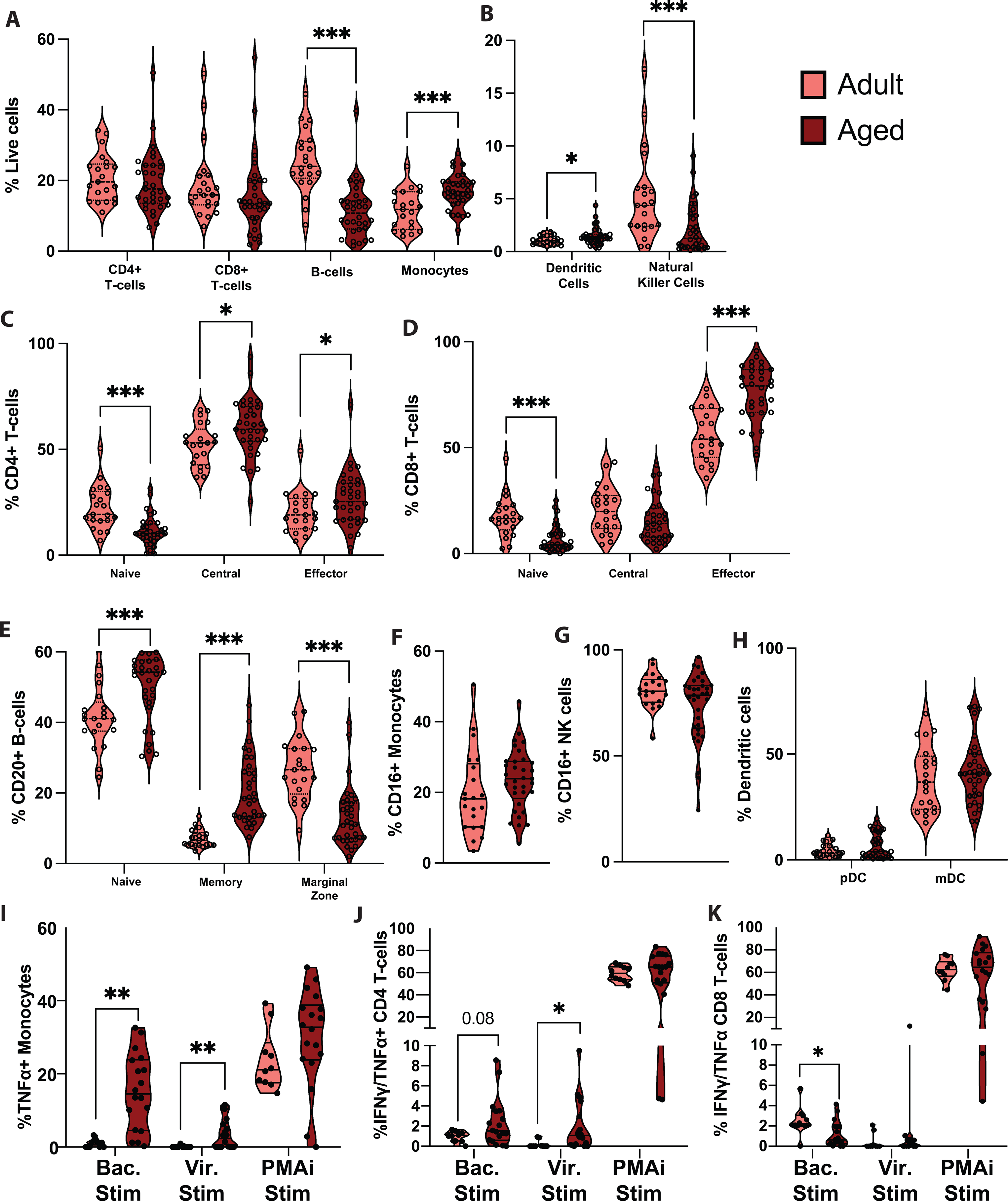
Age-related differences in circulating immune cell frequency and function. Violin plots of immune cell frequency as measured by flow cytometry of BAL cells for (A) major immune cell populations (CD4+ T-cells, CD8+ T-cells, B-cells, Monocytes), (B) less abundant immune cell populations (Dendritic cells, Natural killer cells), (C) CD4+ T-cell subsets, (D) CD8+ T-cell subsets, (E) CD20+ B cell subsets, (F) CD16+ Monocytes, (G) CD16+ Natural killer cells, and (H) Plasmacytoid and Myeloid Dendritic cells. (I) Percent abundance of TNF*α* producing monocytes after 16hr PMAi stimulation. Percent abundance of IFN*γ* and/or TNF*α* producing (J) CD4+ and (K) CD8+ T-cells after 16hr stimulation. Cytokine producing cells were measured using intra-cellular cytokine staining and flow cytometry. Cells were stimulated with either bacterial ligands (Bac. Sim; LPS, FSL-1, Pam3CSK4), viral ligand (Vir. Stim; ssRNA, Imiquimod, ODN2216), or PMAi. Comparisons were made using an unpaired t-test between age-groups for each cell subset (* = p<0.05, ** = p<0.01, *** = p<0.001).

**Figure S2:**
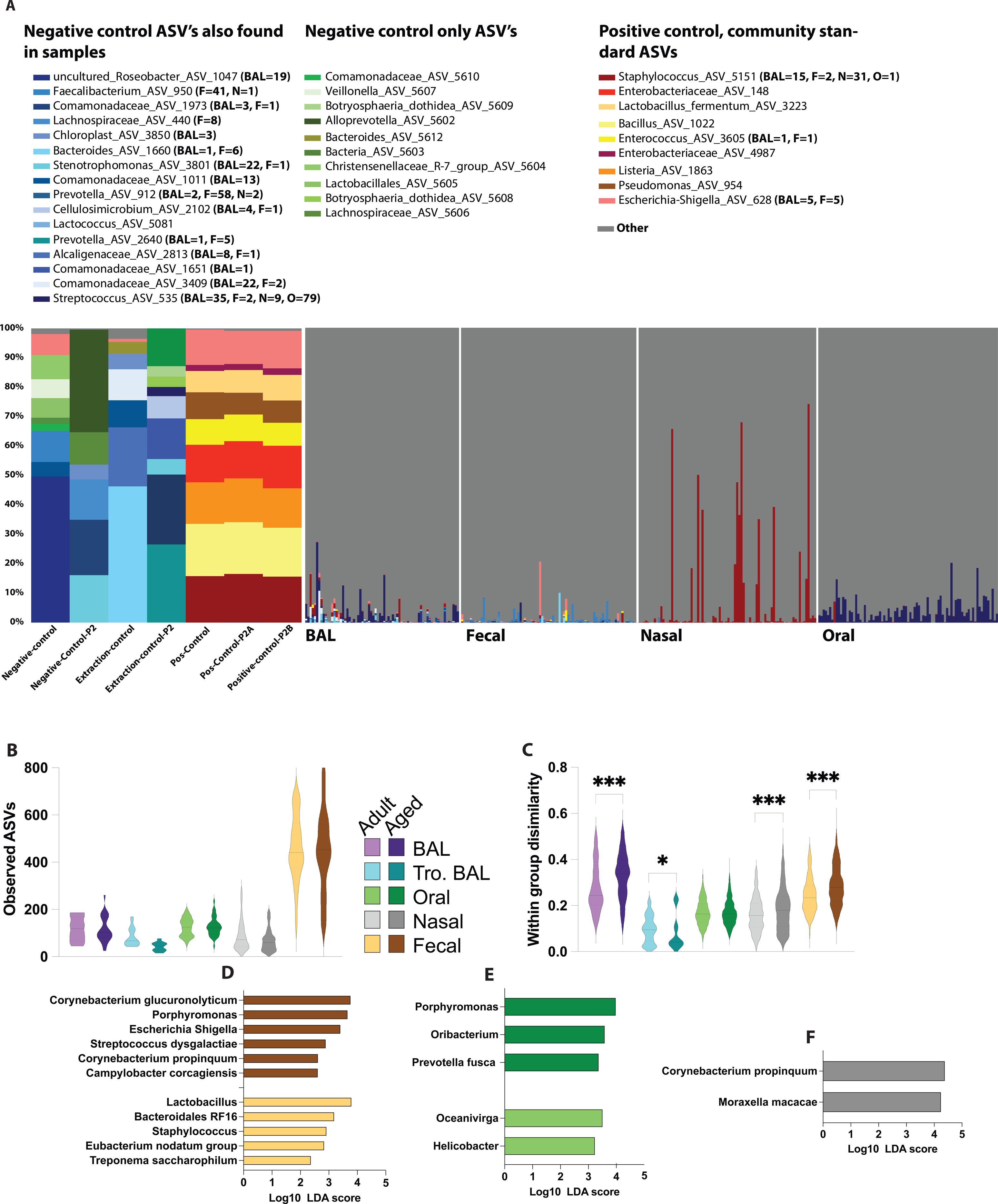
Contribution of environmental contaminates to the lung microbiome and age-related shifts in the rhesus macaque microbiome. (A) Stacked bar plot highlighting taxa found in positive and negative control samples. (B) Violin plots of observed amplicon sequencing variants (ASVs) across sampling sights between age groups. (C) Violin plot illustrating within group weighted UniFrac distances between animals of the same age group within each site. (D-F) Differentially abundant bacterial taxa between age groups at each site (fecal, oral and nasal). Differential abundance was determined using LEFsE (Log10 LDA score > 2) of L7 (species level) taxonomy.

**Figure S3:**
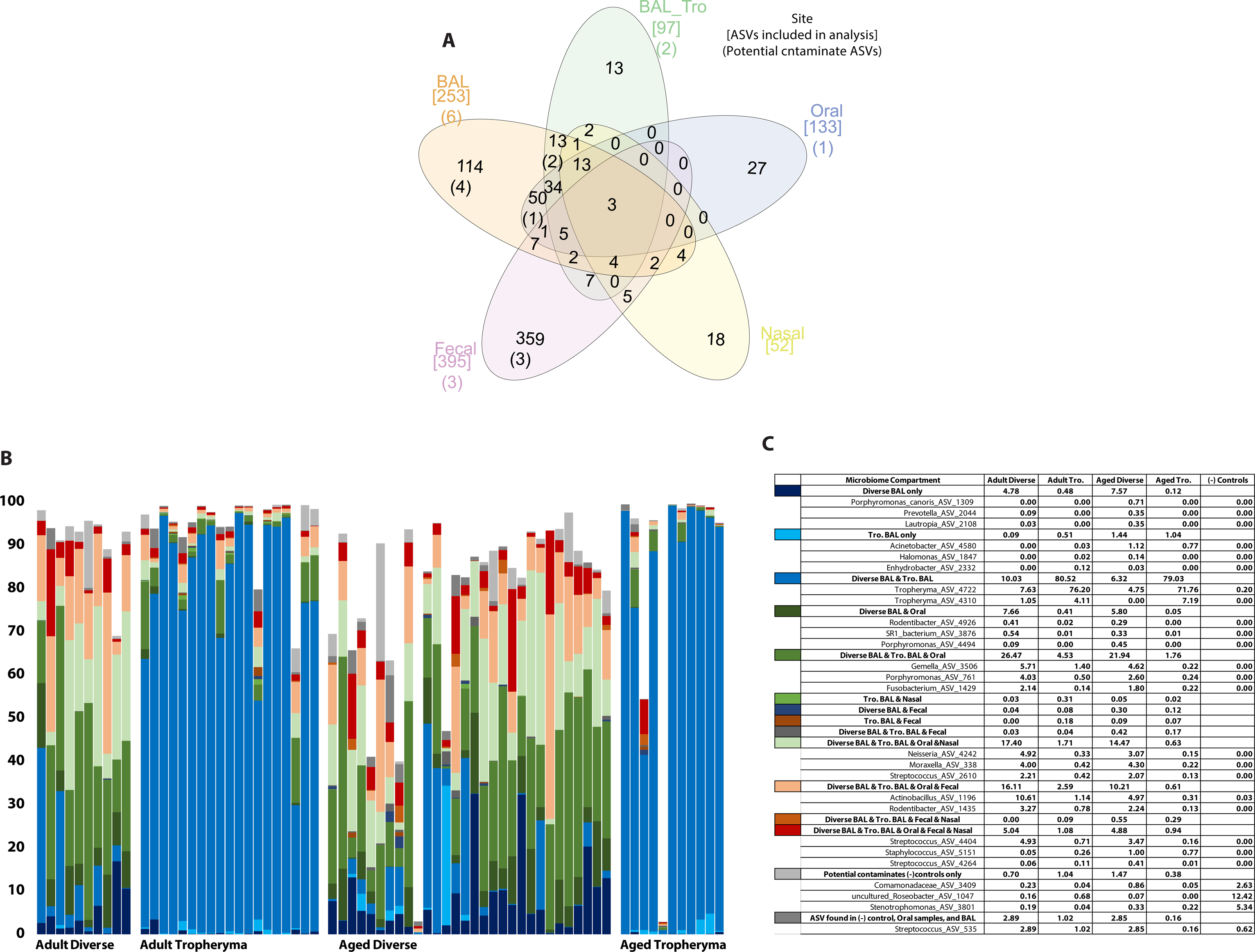
Biogeography of the rhesus macaque microbiome in relation to the lungs. (A) Venn diagram of shared and unique microbial ASV’s found within and between each sampling sites. To be included in this analysis ASV’s had to be detected with >0.01% average relative abundance and present in >10% samples within a site. (B) Stacked bar plot of ASV relative abundance across all samples, colored based on which body sites those ASV’s are shared with. (C) Table with the three most abundant ASVs unique to the lung or shared with other body sites averaged for each age group and BAL microbiome phenotype

**Figure S4:**
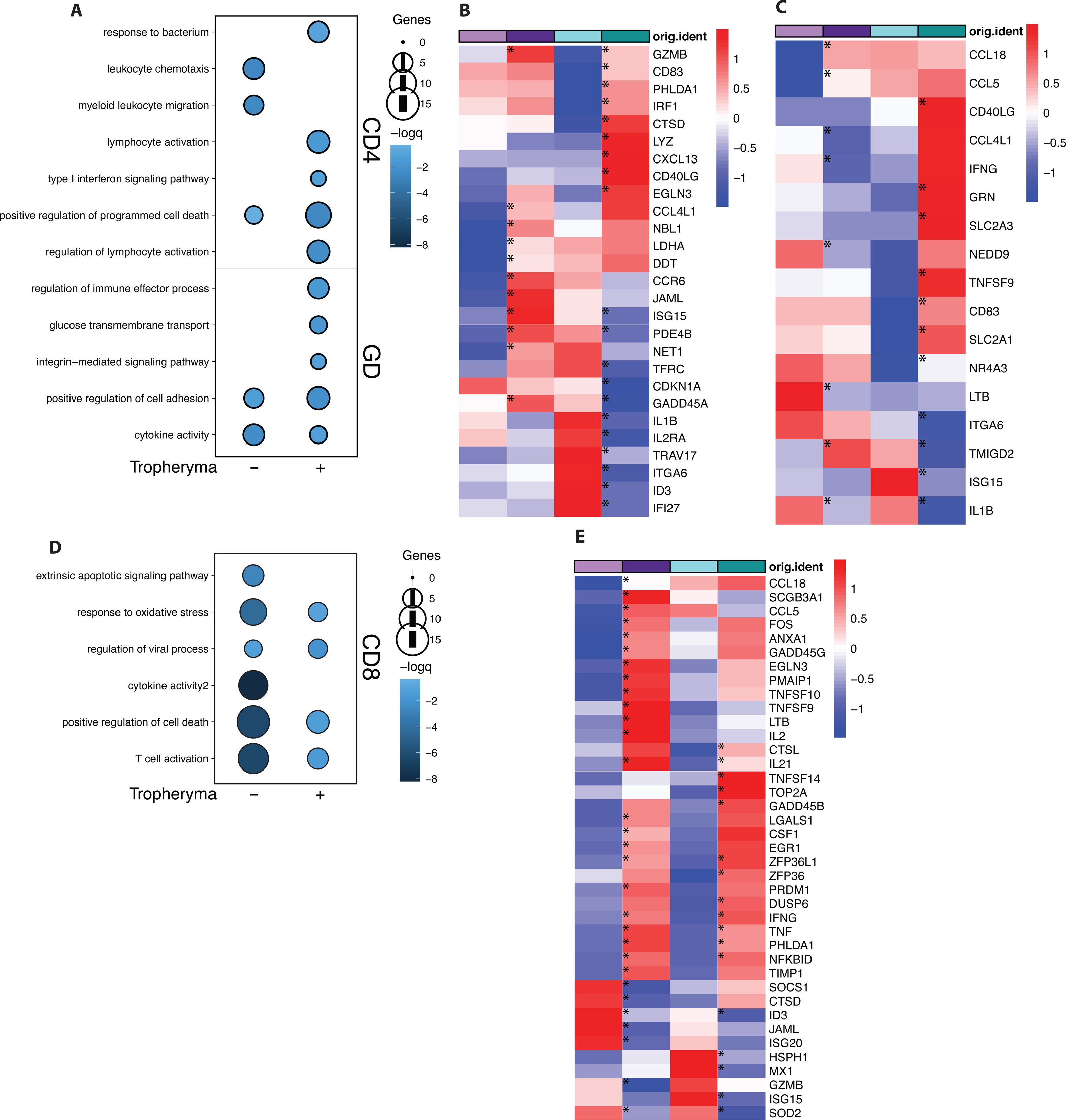
Age- and *Tropheryma*-related shifts in BAL T-cell expression. (A) Functional enrichment of genes differentially expressed in CD4+ T-cells (clusters 0 and 5) and *γδ* T-cells (cluster 2) between adult and aged animals in the absence and presence of *Tropheryma* colonization. Size of the bubble represents statistical significance and number of genes respectively. (B,C) Heatmap of differentially expressed genes within CD4+ T-cells (B) and *γδ* T-cells (C). (D) Functional enrichment of genes differentially expressed in CD8+ T-cells (clusters 1 and 3) between adult and aged animals in the absence and presence of *Tropheryma* colonization. Size of the bubble represents statistical significance and number of genes respectively. (E) Heatmap of differentially expressed genes within CD8+ T-cells. For all heatmaps only genes with p < 0.05 are included; * indicate the comparison (Adult vs. Aged or Adult Tro. vs. Aged Tro) each DEG was derived from.

**Figure S5:**
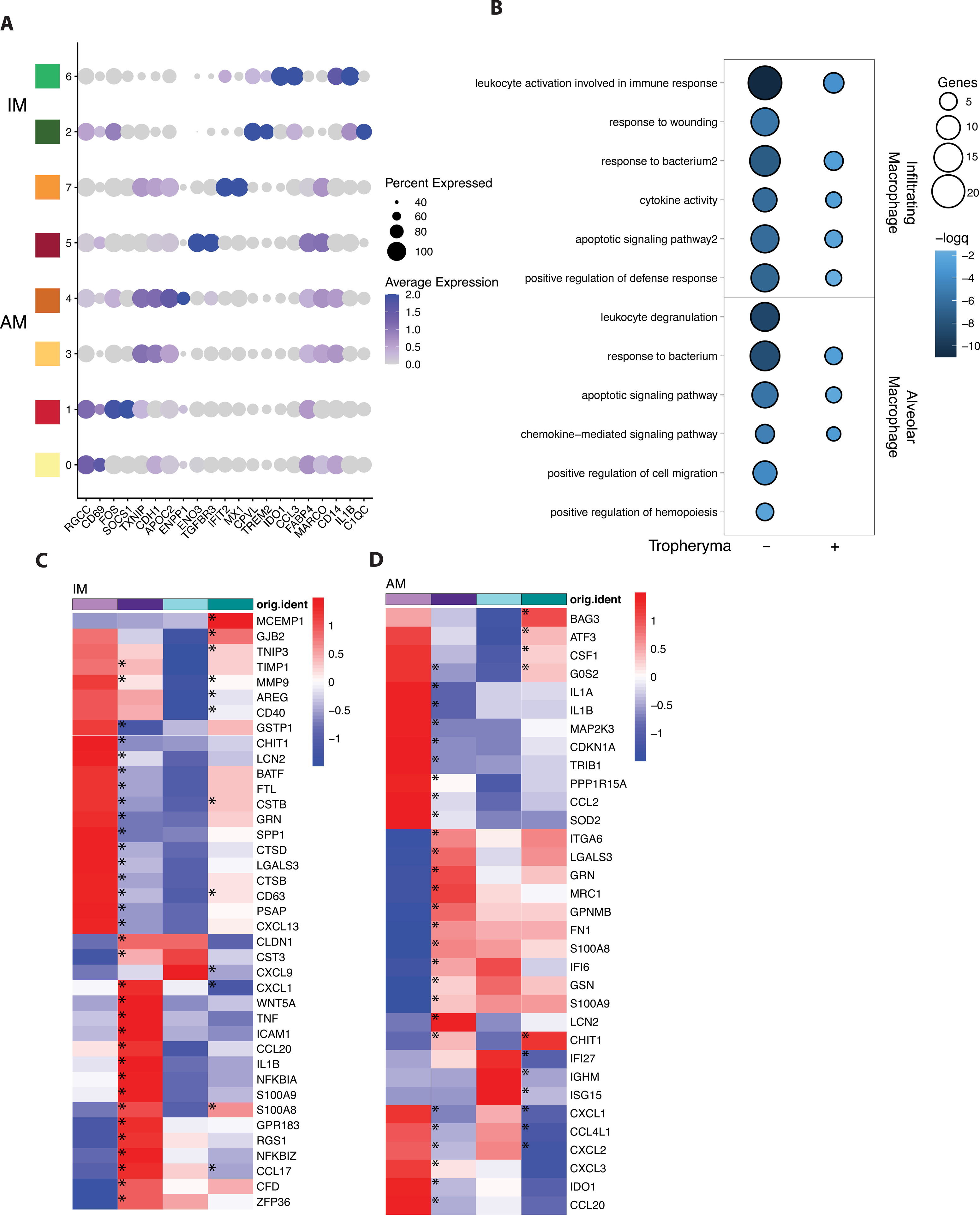
Age and *Tropheryma* related shifts in BAL AM and IM expression. (A) Bubble plot of identifying genes highly expressed in AM and IM cell clusters determined using the FindMarkers function in Seurat. Level of normalized expression is shown using a color scale ranging from low (white) to high (purple). Size of the bubble is representative of the fraction of cells within each cluster expressing the marker. (B) Functional enrichment of genes differentially expressed in IM (clusters 2 and 6) and AM (clusters 0,1,3,4,5,7) between adult and aged animals in the absence and presence of *Tropheryma* colonization. Size of the bubble represents statistical significance and number of genes respectively. (C,D) Heatmap of differentially expressed genes within IM (C) and AM (D). For all heatmaps only genes with p < 0.05 are included, * indicate for which comparisons (Adult vs. Aged or Adult Tro. vs. Aged Tro) each gene was differentially expressed.

**Supplement Table 1:** Cohort information

**Supplement Table 2:** Pulmonary function ANOVA results.

**Supplement Table 3:** Fecal lefse results

**Supplement Table 4:** Single cell cluster markers Figure 3

**Supplement Table 5:** Single cell cluster markers Figure 4

**Supplement Table 6:** Single cell cluster markers Figure 5

## METHODS

### Cohort and sample collection

Samples were obtained from a total of 82 rhesus macaques and a full breakdown of animal data and experiment can be found in supplemental table 1 (**Supp. Table 1**). All rhesus macaque studies were overseen and approved by the OHSU/ONPRC and the NIA Intramural Institutional Animal Care and Use Committees (IACUC) per the National Institutes of Health guide for the care and use of laboratory animals. Animals were housed per the standards established by the US Federal Animal Welfare Act and The Guide for the Care and Use of Laboratory Animals. Animals observed at ONPRC were pre-screened to establish their suitability for experimental procedures. All animals 18 years and older receive physical exams every six months, and are also tested for normality in CBC, blood chemistry and a fecal occult test. An electronic animal records system documents an animals long-term health record and identifies any criteria that invalidates its usage. Whole blood samples were collected in EDTA vacutainer tubes. Peripheral blood mononuclear cells (PBMC) were obtained by standard density gradient centrifugation over the blood separation polymer Ficoll in Sepmate tubes. BAL cells were isolated from total BAL by centrifugation at 1200 rcf for 15 minutes. PBMC and BAL cells were cryo-preserved using 10% DMSO/FBS and Mr. Frosty Freezing containers (Thermo Fisher Scientific) at −80C then transferred to a cryogenic unit until analysis. Whole BAL for microbiome analysis was collected prior to the isolation of BAL cells. All swabs, (nasal, oral, fecal) were collected at time of BAL.

### Pulmonary function testing

For pulmonary function testing, rhesus macaques were anesthetized with ketamine and dexmedetomidine then intubated. Pulmonary function tests were performed using the Buxco pulmonary function apparatus from DSI/Harvard Bioscience as previously described (60, 61). Gas exchange (DLCO) and alveolar volume were measured using carbon monoxide (CO) diffusion after the methods of Castillo et al (104) and Fallica et al (105). The reference gas containing 0.5% CO , 0.5% neon, 21% O2, 78% N2 was injected into the lungs, withdrawn after 4 seconds and gas levels measured by gas chromatography. All tests were performed a minimum of 3 times. After testing was complete, animals were extubated, monitored and returned to their home cages. Thoracic CT scanning with controlled lung inflation was performed in a subset of animals (5 young, 6 old) on a GE Optima CT 660 with a 64-slice detector and analyzed by a cardiothoracic radiologist.

### 16S amplicon sequencing

Total DNA was extracted from 400ul of BAL fluid or swabs (fecal/oral/nasal) using the DNeasy Powersoil Pro Kit (Qiagen, Valencia, CA, USA). The hypervariable V4-V5 region of the 16S rRNA gene was amplified using PCR primers (515F/806R with the forward primers including a 12-bp barcode) (*70*). PCR reactions were conducted in triplicates and contained 12.5 ul GoTaq master mix, 9.5 ul nuclease-free H20, 1 ul template DNA, and 1 ul 10uM primer mix. Thermal cycling parameters were 94°C for 5 minutes, 35 cycles of 94°C for 20 seconds, 50°C for 20 seconds, 72°C for 30 seconds, followed by 72°C for 5 minutes. PCR products were purified using a MinElute 96 UF PCR Purification Kit (Qiagen, Valencia, CA, USA). Libraries were sequenced (2 x 300 bases) using an Illumina MiSeq.

Raw FASTQ 16S rRNA gene amplicon sequences were uploaded and processed using the QIIME2 analysis pipeline (106). Briefly, sequences were demultiplexed and the quality filtered using DADA2 (107), which filters chimeric sequences and generates an amplicon sequence variant (ASV) table equivalent to an operational taxonomic unit (OTU) table at 100% sequence similarity. Sequence variants were then aligned using MAFFT (108) and a phylogenetic tree was constructed using FastTree2 (109). Taxonomy was assigned to sequence variants using q2-feature-classifier against the SILVA database (release 138) (110). To prevent sequencing depth bias samples were rarified to 20,000 sequences per sample before alpha and beta diversity analysis. QIIME 2 was also used to generate the following alpha diversity metrics: richness (as observed ASV), Shannon evenness, and phylogenetic diversity. Beta diversity was estimated in QIIME 2 using weighted and unweighted UniFrac distances (111).

### Full length 16S sequencing and tree building

For *Tropheryma* dominated animals the full length 16S rRNA gene was amplified using the B8F-1492R primers. Cleaned PCR reactions were then sequenced bi-directionally using Sanger di-deoxy sequencing. Paired sequences were then merged, trimmed, and aligned against known *Tropheryma* 16S rRNA gene sequences using the DNAsubway pipeline (https://dnasubway.cyverse.org). The final alignment was exported as a newick file and plotted in iTOL (itol.embl.de).

### Flow cytometry

To profile adaptive immune cell subsets 1 × 10^6^ BAL cells or PBMC were stained using antibodies against CD4, CD8b, CD95, CD28, CD20, CD27 and IgD to delineate naïve and memory CD4 and CD8 T cells, as well as CD20 B cell subsets. T cells were then divided into Naive (CD28+, CD95−; Naive), central/transitional memory (CD28+ CD95+; CM), and Effector memory (CD28− CD95+; EM). A second tube of 1 × 106 BAL cells or PBMC was stained using antibodies against: CD3, CD20, HLA-DR, CD14, CD11c, CD123, CD16, CD8a, and CD206 (BAL only) to delineate Alveolar macrophages (CD3− CD20− CD206+) Infiltrating macrophages (CD3− CD20− CD206-CD14 + HLA-DR+), dendritic cells (DC, CD3− CD20-CD14− HLA-DR+), and natural killer (NK, CD3-CD20-CD14− HLA-DR− CD8a+) cell subsets. Macrophages were further subdivided into classical (CD16−) and non-classical macrophages (CD16+). DCs were further subdivided into myeloid DC (mDC, CD123− CD11c+) and plasmacytoid DC (pDC, CD123+CD11c−). All flow cytometry samples were acquired using Attune NxT (Life Technologies, Carlsbad, CA, United States) and analyzed using FlowJo (TreeStar, Ashland, OR, United States).

### Stimulation and ICS

BAL cells (5 × 10^5^/well) were thawed and incubated for 16 h in the absence (unstimulated) or presence (stimulated) of 5 ng/ml PMA and 1 μg/ml Ionomycin (Sigma-Aldrich, St. Louis, MO, United States). PBMC cells (5 × 105/well) were thawed and incubated for 16 h in the absence (unstimulated) or presence (stimulated) of PMAi (5 ng/ml PMA and 1 μg/ml Ionomycin) or Bacterial TLR antigen cocktail (LPS, FSL-1, Pam3CSK4) or Viral TLR antigen cocktail (ssRNA, Imiquimod, ODN2216). All cells were incubated with brefeldin A at 37°C in a humidified incubator (5% CO2). After incubation cell were stained using antibodies against CD4, CD8b, CD28, CD20, CD14 and HLA-DR to delineate total T-cell and monocyte populations. Cells were next fixed and permeabilized and stained with IFN*γ*, TNF*α* and IL-6.

### Single cell RNA library preparation

Cryopreserved BAL cells from a subset of animals (n=3/group for Adult No Tro., Adult Tro., Aged No Tro., and Aged Tro.) were thawed. Samples were washed three times in ice cold PBS supplemented with 2% FBS and sorted on the FACS Aria Fusion (BD Biosciences) with Ghost Dye Red 710 (Tonbo Biosciences) for dead cell exclusion. Live cells were counted in triplicates on a TC20 Automated Cell Counter (BioRad) and pooled in groups of 3 based on host age and *Tropheryma* status. Pooled cells were resuspended in ice cold PBS with 0.04% BSA in a final concentration of 1200 cells/uL. Single cell suspensions were then immediately loaded on the 10X Genomics Chromium Controller with a loading target of 17,600 cells. Libraries were generated per manufacturer’s recommendation Chromium Single Cell 3’ Feature Barcoding Library Kit using the v3.1 chemistry (10X Genomics, Pleasanton CA). Libraries were sequenced on Illumina NovaSeq with a sequencing target of 30,000 reads per cell RNA library and 2,000 reads per cell HTO barcode library.

### Single cell RNA-Seq data analysis

Raw reads were aligned and quantified using the Cell Ranger Single-Cell Software Suite (version 4.0, 10X Genomics) against the Mmul_8 rhesus macaque reference genome using the STAR aligner. Downstream processing of aligned reads was performed using Seurat (version 4.0). Droplets with ambient RNA (cells fewer than 200 detected genes) and dying cells (cells with more than 25% total mitochondrial gene expression) were excluded during initial QC. Data objects from all groups were integrated using *Seurat* (112). Data normalization and variance stabilization was performed on the integrated object using the *SCTransform* function where a regularized negative binomial regression corrected for differential effects of mitochondrial cell cycle gene expression levels. The R package doublet finder (113) along with manual curation was used to remove suspected doublet cells based on the expression of multiple lineage marker. Dimensional reduction was performed using *RunPCA* function to obtain the first 30 principal components followed by clustering using the *FindClusters* function in Seurat. Clusters were visualized using the UMAP algorithm as implemented by Seurat’s *runUMAP*function. Cell types were assigned to individual clusters using *FindMarkers* function with a fold change cutoff of at least 0.4. Clusters from two major cell types (T/NK cells and Macrophages) were further subsetted from the total cells and re-clustered to identify minor subsets within those groups.

### Statistical analysis

Differential expression analysis was performed using MAST using default settings in Seurat. All disease comparisons were performed relative to healthy donors from corresponding age groups. Only statistically significant genes (Fold change cutoff ≥ 1.5; adjusted p-value ≤ 0.05) were included in downstream analysis. Significance of two group comparisons (i.e. all Adult vs. all Aged) were measured using the non-parametric Mann-Whitney U-test. Significance of 3+ group comparison was measured using the non-parametric Kruskal-Wallis ANOVA with Dunn’s multiple comparison test for post-hoc pairwise comparison when the initial ANOVA was significant. The LEfSe algorithm was used to identify differentially abundant taxa and pathways between groups with a logarithmic Linear discriminant analysis (LDA) score cutoff of 2 (114).

